# Neuralized-like proteins differentially activate Notch ligands

**DOI:** 10.1101/2024.09.20.614084

**Authors:** Alina Airich, Oren Gozlan, Ekaterina Seib, Lena-Sophie Wilschrey, Gittel Leah Shaingarten, David Sprinzak, Thomas Klein

**Author notes:** Correspondence: David Sprinzak, Thomas Klein, **Email:**. Equal contributions (Alina Airich, Oren Gozlan).

## Abstract

Notch signalling is a major signalling pathway coordinating cellular processes between neighbouring animal cells. In Drosophila, two ubiquitin ligases, Neuralized (Neur) and Mindbomb1 (Mib1), regulate Notch ligand activation and are essential for development. However, the mammalian orthologs of Neur, Neuralized-like (NEURL) 1 and 2, do not appear to be crucial for development, as double knock-out mice show no developmental defects. Thus, it is unclear if and how NEURL proteins regulate the four mammalian Notch ligands. To address these questions, we examined NEURL proteins’ ability to activate Notch ligands in humanized *Drosophila* and mammalian cell culture. We found that, unlike MIB1, NEURL proteins activate Notch only with a subset of mammalian ligands, which contain a Neuralized binding motif. This motif has the consensus sequence NxxN, present only in Notch ligands DLL1 and JAG1, but not in DLL4 and JAG2. Overall, we show that NEURL proteins activate specific Notch-ligands, suggesting a differential regulatory mechanism of Notch activation in mammals, which can potentially explain the limited role of NEURL proteins in mammalian development and homeostasis.

## Introduction

Notch signalling is a highly conserved pathway that coordinates cellular processes between neighbouring cells throughout the animal kingdom^1–3^. Notch signalling relies on the interaction between membrane-bound Delta-Serrate-Lag2 (DSL) ligands on a sender cell and Notch receptors on a receiver cell. While highly conserved, there are differences between the number and type of Notch ligands and receptors across animal species. In *Drosophila*, there are two ligands, termed Delta (Dl) and Serrate (Ser), and one Notch receptor. In mammals, there are five ligands, three from the Delta-like (Dll) family (Dll1, Dll3, Dll4), and two from the Jagged (Jag) family (Jag1 and Jag2). There are four mammalian Notch receptors (Notch1-4). All ligands and receptors can interact with each other, albeit with different signalling strengths^1,3^. Yet, it is unclear how different receptor-ligand interactions are specifically regulated to induce different cellular outcomes.

Canonical Notch signalling is initiated by the binding of DSL ligands in the sender cell to the Notch receptor in the receiver cell (Fig.1A). The binding results in a conformational change in Notch, which leads to the cleavage of the receptor - first by ADAM10 (S2-cleavage) and then by the 𝛾-secretase complex (S3-cleavage), releasing the intracellular domain of Notch (NICD) into the cytosol. From there, the NICD translocates into the nucleus, where it regulates the expression of target genes.

**FIG. 1.**
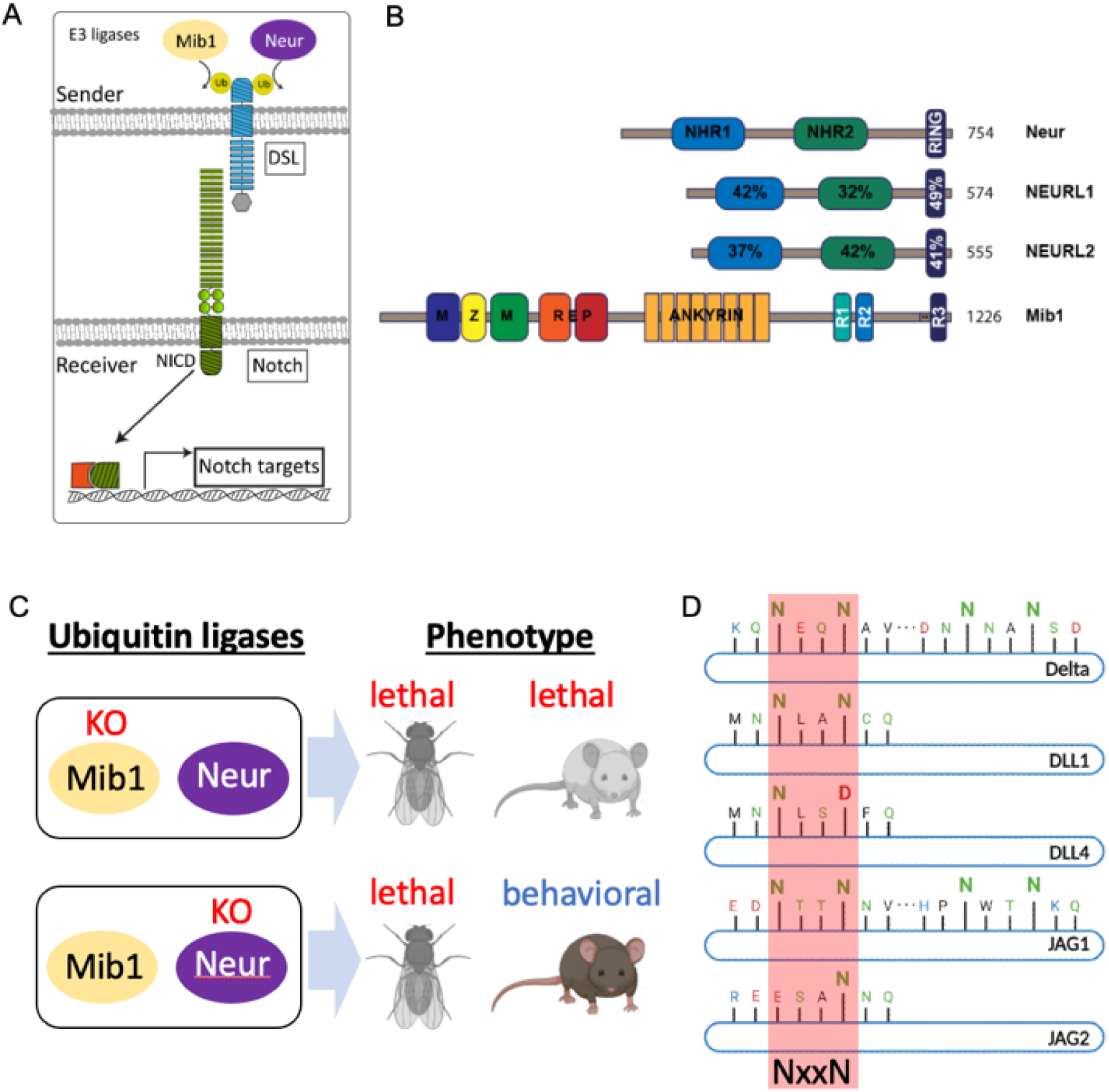
The role of Neur and Neurl’s in ligand-dependent activation of the Notch pathway. (A) E3-ligases Mib1 and Neur mediate ubi of the ICDs of the DSL ligands in the sender cell, which in turn leads to the full activation of the Notch receptors in the receiver cell. (B) The structure of Neur, NEURL-1, NEURL-2 and MIB1. Neur family members have two NHR-domains, followed by a RING domain. MIB1 consists of the MZM and REP domains, followed by ANK repeats and three RING domains. (C) The distinct KO-phenotypes of MIB1 and Neur in Drosophila and mouse. (D) The position of the NxxN consensus sequence in the ICDs of the mammalian ligands. It is recognisable in Dl, DLL1, and JAG1. DLL4 and JAG2 appear to have a cryptic sequence where one N is replaced by E or Q. The sequences are conserved among mammalian orthologs (see Fig. 1 S1A).

In *Drosophila*, two E3-ligases, termed Neuralized (Neur) and Mindbomb1 (Mib1), are involved in the activation of Notch ligands in the sender cells^4,5^ (Fig. 1B). Both types of ligases share no overall sequence or structural similarity beyond the RING domains at their C-terminus (Fig. 1B). Nevertheless, both induce endocytosis of the ligands either by ubiquitylation (ubi) of the ligand’s lysine residues found on their intracellular domain (ICD) or, in the case of Neur, in a ubi-independent manner^6^. For their activation, the two E3-ligases bind the ligands at different epitopes. Neur binds its substrates at the Neuralized-binding motif (NBM) with the consensus sequence NxxN, located in the ligands’ ICD^7^ This NxxN sequence is found in all Neur substrates in *Drosophila,* including Ser, Dl, and the Bearded proteins. After binding and ubiquitylation, the ligands are endocytosed, generating a pulling force that induces a conformational change in Notch receptor, which is required for ADAM10-dependent S2-cleavage of the receptor^8^.

In mammalian genomes, one Mib1 ortholog and two Neur orthologs, Neuralized-like-1 (Neurl-1) and Neuralized-like-2 (Neurl-2), have been identified. A knock-out (KO) of Mib1 in mice is lethal and causes the expected Notch-like developmental phenotypes^9^ (Fig. 1C). In contrast, Neurl-1 and −2 double KO mice survive to adulthood without any obvious developmental defects^10^ (Fig. 1C). This has led to the general belief that only Mib1, but not Neurl-1 and −2, regulates Notch ligand activity in mammals. Human MIB1 was shown to bind in a bipartite mode at different epitopes of the ligands’ ICD^11,12^.

More recently, analysis of the Neurl-1 and −2 double KO mice revealed that their function is largely restricted to postnatal cognitive processes, such as synaptic plasticity and spatial memory^13,14^. Interestingly, conditional KO of Jag1 and Notch1 in the brain has also been associated with spatial memory^13,15,16^. Given the similarities of these phenotypes, this may suggest that Neurl proteins have a role in the Notch pathway that is more restricted to specific neuronal processes in adults. Thus, it is unclear why in *Drosophila*, both Mib1 and Neur are essential for development, while only Mib1, but not Neurl proteins, is crucial for mammalian development.

Here, we investigated the ability of human NEURL-1 and NEURL-2 to activate the mammalian Notch ligands in both, humanized *Drosophila* and mammalian cell culture. We found that both NEURL proteins can activate a subset of the ligands. The ability of NEURL proteins to activate ligands depends on the presence of the consensus sequence NxxN motif in their ICD, which we show here is a NBM also in mammalian ligands. Ligands that contain this NBM in their ICD, DLL1 and JAG1, can activate Notch receptors in a NEURL-dependent manner both *in vivo* (*Drosophila*) and *in vitro* (cell culture assay). Ligands lacking the NBM, DLL4 and JAG2, cannot activate NEURL-mediated Notch signalling in either system. Moreover, mutating the NBM in the ICD of DLL1 and JAG1 abolishes NEURL-mediated Notch activity but not MIB1-mediated activity. Introducing an NBM in the ICD of DLL4 rescued NEURL activity in both *Drosophila* and mammalian cell culture. Moreover, we found increased co-localization of NEURL-1 and Notch ligands only in ligands activated by NEURL-1 (DLL1 and JAG1), suggesting stronger interaction of NEURL-1 with these ligands. Thus, our results indicate that NEURL-1 and NEURL-2 can differentially activate ligands containing the NBM (DLL1 and JAG1), while showing no activation of the other ligands (DLL4 and JAG2). We suggest that the cognitive phenotypes associated with Neurl-1 and −2 in mice may be due to Notch-dependent processes restricted to specific ligands.

## Results

A sequence comparison of Dl with the mammalian DSL-ligands revealed that the ICDs of Dll1 and Jag1 in several vertebrate species contain a sequence at the N-terminus that matches the NxxN consensus motif of a NBM (NLAN in DLL1 and NTTN in JAG1 in humans, Fig. 1D, Fig. 1S1A). The strong conservation suggests that the NxxN motif undergoes evolutionary selection, indicating its potential importance for the function of the ligands. While the DLL4 ICD contains a sequence similar to the DLL1 NBM, it does not have a complete NxxN motif, as the second N is replaced by a negatively charged aspartic acid (D) (NLS**D,** Fig. 1D, Fig. 1S1A). Similarly, in the case of JAG-2, a sequence showing similarity to the NxxN motif exists, where one N is replaced by a negatively charged glutamic acid (E) (ESAN) (Fig. 1D, Fig. 1S1A).

### Neurl-1 and −2 can activate the Notch pathway in *Drosophila*

To determine whether Neurl proteins function as homologs of Neur and can activate specific ligands, we analysed their activity in complementary experiments using *Drosophila* and mammalian cell culture. For reasons of availability, we used mouse Neurl-1 and human NEURL-2 for our *Drosophila* experiments. Mouse Neurl-1 is 94.3% identical to human NEURL-1^17^. *Drosophila* experiments showed Neurl activity in *in-vivo* setting, while the cell culture assays showed the same results in quantitative *in-vitro* mammalian setting.

We first asked whether Neurl-1 and NEURL-2 can activate the endogenous ligands of *Drosophila*. We therefore, expressed Neurl-1-HA and NEURL-2-HA in *mib1* mutant wing imaginal discs using *ci*Gal4, which drives expression of UAS constructs throughout the anterior compartment (Fig. 2A). In this genetic context, the expressed Neur and NEURL-variants are the only E3-ligases present in the discs that can activate the Notch pathway. The constructs used here are inserted into the same genomic landing site to ensure comparable expression levels. The loss of *mib1* function results in a strong loss of Notch activity leading to the loss of *wingless* (Wg) expression along the dorso-ventral compartment boundary (D/V-boundary) and a dramatic size reduction of the wing primordium^18,19^ (Fig. 2A’, compare with B’). The expression of Wg depends on the activity of both ligands, with Ser being the more important one^20,21^. We found that the expression of the Notch target gene Wg along the D/V-boundary is re-established in the region of expression of the NEURL proteins (Fig. 2A’’-A’’’’, compare with A’ and B’). There are differences in the degree of rescue of Notch activity between the two NEURL proteins compared to Neur. Both NEURL proteins exhibit weaker activation compared to Neur, with Neurl-1 showing better rescue than NEURL-2 (length of DV stripe in Fig. 2A’’-A’’’’). Hence, the results indicate that each of the two NEURL proteins can activate endogenous *Drosophila* ligands to a level sufficient to induce the expression of Wg.

**FIG. 2.**
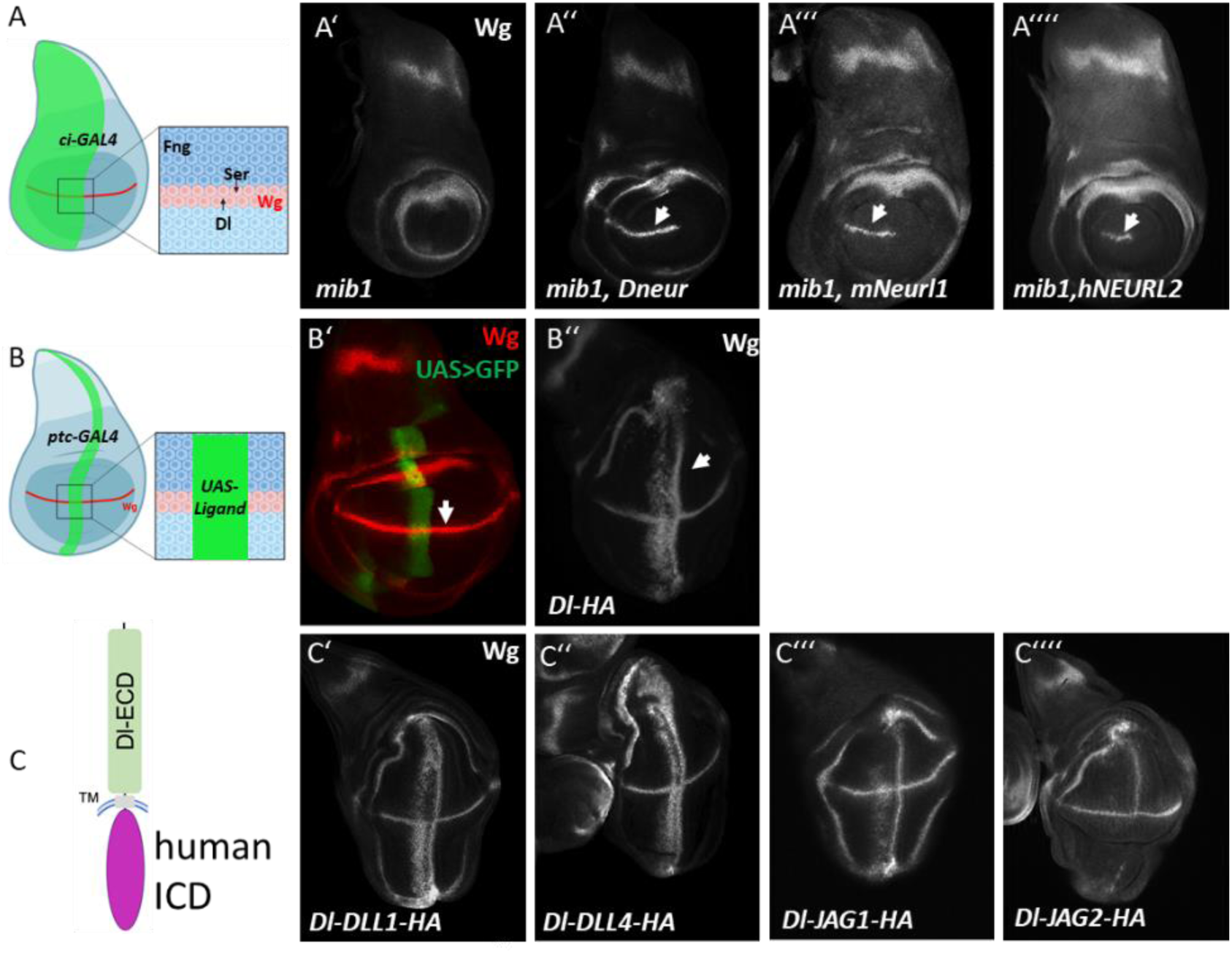
Activation of the human ligands by Neur, NEURL-1 and NEURL-2 in the wing imaginal disc in Drosophila. (A) Cartoon of the expression domain of Ci-FGal4, which is expressed throughout the anterior compartment (green area). At the D/V boundary, interaction between dorsal and ventral boundary cells, mediated by Dl and Ser activate the Notch pathway and induce the expression of target genes, such as Wg. (A’) In mib1 mutant discs, the expression of Wg along the D/V boundary is lost. (A’’-A’’’’) Expression of Neur, Neurl1 and Neurl2 in mib1 mutant wing discs results in the re-establishment of Wg expression along the D/V boundary (arrow in A’’-A’’’’). (B, B’) Expression of ptcGal4 in the wing imaginal disc. (B) Cartoon of the expression. (B’) ptc-Gal4 is expressed in a broad stripe at the anterior side of the A/P compartment boundary. The expression domain runs perpendicular to the expression of Wg along the D/V boundary (arrow). (B’’) Expression of Dl with ptc-Gal4 results in the induction of two ectopic stripes of Wg expression (arrow). (C) Cartoon of the hybrid ligands generated for this study. In these hybrids, the ICD of Dl is replaced by that of the mammalian ligands. (C’-C’’’’) Expression of Dl-DLL1, Dl-DLL4, Dl-JAG1 and Dl-JAG2 with ptc-Gal4 results in the ectopic expression of Wg.

### The ICDs of the human ligands can mediate the Mib1-dependent activation of Dl-hybrids in *Drosophila*

To rigorously test the activity of the ICDs of mammalian ligands in *Drosophila*, we generated hybrid ligands in which the ICD of Dl is replaced by that of human ligands (Dl-DLL1, Dl-DLL4, Dl-JAG1, and Dl-JAG2, Fig. 2C). We excluded the ICD of DLL3 since it is not a ligand capable of trans-activation of Notch^22,23^. The constructs were inserted into the same genomic landing-site and expressed with the Gal4-system.

We first expressed the Dl-hybrids in wildtype discs using *ptc*Gal4, which drives expression visible as a stripe along the anterior side of the anterior-posterior compartment boundary (A/P-boundary, Fig. 2B, B’). Expression of Dl with *ptc*Gal4 induces the expression of Notch target genes, including Wg, in two ectopic stripes running perpendicular to the endogenous expression domain that straddles the D/V-boundary (Fig. 2B’, B’’). Ectopic expression of all four Dl-hybrids with *ptc*Gal4 resulted in similar ectopic activation of the Notch pathway in the wing disc, as indicated by the induction of ectopic expression of Wg (Fig. 2C’-C’’’’). While the expression of Dl-DLL1, Dl-DLL4, and Dl-JAG1 resulted in the Notch-pathway activation at a level comparable to Dl, there was less ectopic expression of ectopic Wg in the case of Dl-JAG2 (Fig. 2C’-C’’’, compare with B’’). None of the hybrid ligands were able to activate the Notch pathway in *mib1* mutant discs (Fig. S1B-E). Since Mib1 is the major E3 ligase broadly expressed in the wing disc at this stage, these results reveal that the hybrid ligands can induce Mib1-mediated Notch activation in *Drosophila*^18,19,24^.

### Neurl-1 and Neurl-2 differentially activate hybrid DSL-ligands in humanized Drosophila

To test the ability of Neur and its mammalian orthologs to activate our Dl-hybrid constructs, we co-expressed the hybrid ligands with the Neurl’s in *mib1* mutant discs. In this genetic situation, the Neur proteins are the only E3 ligase able to activate the Notch ligands. Consequently, we co-expressed Dl-hybrids in *mib1* mutant discs in various combinations. Co-expression of Dl with Neur in *mib1* mutants results in the induction of two ectopic stripes of Wg^6^. We found that activation of Notch was observed also upon co-expression of either Neur, Neurl-1 or NEURL-2 with Dl-DLL1 or Dl-JAG1, but not with Dl-DLL4 or Dl-JAG2 (Fig. 3A-C’’’’). This matched our sequence comparison result that showed that DLL1 and JAG1, but not DLL4 and JAG2, contain the NxxN motif. The activation of Dl-JAG1and Dl-DLL1 induced by Neurl-1 was stronger than that induced by NEURL-2 (Fig. 3B-C’’’’). This suggests that Neurl-1 is a better activator of DSL-ligands than NEURL-2. We found that also a Dl-hybrid containing the ICD of rat Dll1 (Dl-rDll1), which contains the conserved NxxN sequence, could be activated by all three Neur proteins and Mib1 in a manner indistinguishable from Dl-DLL1 (human DLL1, Fig. 3S1A-A’’’). In addition, it could also be activated by Neurl-2 (Fig. 3S1A’, A’’’’). This confirms the broad activation of Dll1 family members by the NEURL proteins. Overall, our findings demonstrate that Neur and its mammalian orthologs selectively activate Notch ligands, highlighting their differential regulatory roles, and confirm that this differential activation occurs also in our hybrid humanized drosophila ligands.

**FIG. 3.**
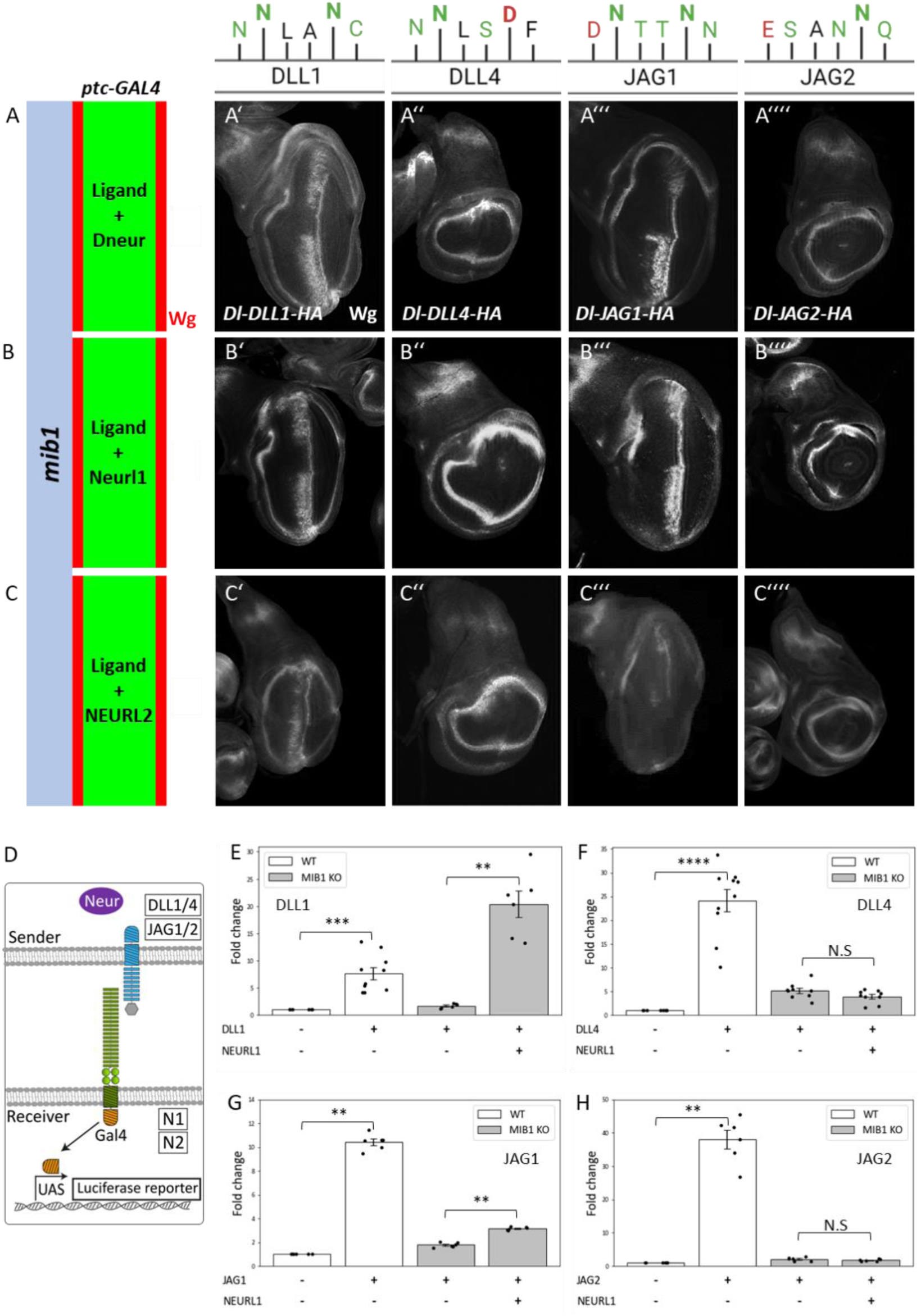
Activation of the mammalian ICDs of DSL ligands by Neur, Neurl1 and Neurl2. (A-C’’’’) Co-expression of ligand and Neur variants in mib1 mutant discs. Only Dl-DLL1 and Dl-JAG1, which contain a NxxN sequence can be activated by Neur, Neurl1 and Neurl2. (D-I) Mammalian cell culture assay for activation of Notch ligands by NEURL-1. Luciferase activity assay results for the different Notch ligands with or without NEURL-1 in a MIB1 KO background. DLL1/4 and their variants were co-cultured with NOTCH1-Gal4 while JAG1 and its variant were co-cultured with NOTCH2-Gal4. 24h after being co-cultured, the cells were lysed and their Notch activity was measured. For each plot the results represent the fold change compared to the negative control. P-values were measured by Mann-Whitney test (* - p<0.05, ** - p<0.01, *** - p<0.001, n>=3).

### NEURL-1 differentially activates DSL-ligands in mammalian cells

We next wanted to test whether NEURL-1 differentially activates DSL-ligands in mammalian cell culture. To assess NEURL-1 activity, we performed a Notch activity assay by co-culturing sender cells expressing full length human DSL ligands (DLL1, DLL4, JAG1, JAG2) and human NEURL-1 with receiver cells expressing a Notch reporter (Fig. 3D-H)^25^. We used human bone osteosarcoma cells (U2OS) from which MIB1 was knocked out as sender cells (MIB1-KO cells)^26^. As receiver cells, we used U2OS cells stably expressing hybrid NOTCH-Gal4 receptors, and transfected them with a UAS-Luciferase reporter gene^25,27^. Notch activity in the receiver cells leads to the release of the intracellular Gal4, which activates the reporter gene (Fig. 3D). Since JAG1 is a very weak activator of Notch1 receptors^28^, we used NOTCH1-Gal4 receiver cells to test DLL1 and DLL4 activity and NOTCH2-Gal4 receiver cells to test JAG1 and JAG2 activity^27^. All ligands were able to activate Notch when expressed in wildtype (WT) U2OS cells that contain endogenous MIB1 (Fig. 3E-H, white bars). MIB1-KO abolished the activity of all ligands (Fig. 3E-H, grey bars), showing that MIB1 is the predominant E3 ligase in our cells and that it is required for Notch ligand activity. When NEURL-1 was co-expressed with Notch ligands in MIB1-KO cells, we observed a rescue of Notch activity only with sender cells expressing DLL1 and JAG1, but not with DLL4 and JAG2 (Fig. 3E-H, grey bars). These results indicate that only ligands containing the NxxN motif could be activated by NEURL-1, in agreement with the *Drosophila* experiments and the prediction from our sequence analysis.

We attempted to test the ability of NEURL-2 to activate Notch ligands in cell culture. However, we found that over-expression of NEURL-2 was toxic to the cells and could not be maintained for the extended time periods required for the assay. Nevertheless, Neurl-2 was able to activate rDll1 and JAG1, but not DLL4 and JAG2 in our *Drosophila* assay (Fig. 3A-C’’’’, Fig. 3S1A, A’’’’).

### The NxxN motif is required and sufficient for activation of Notch ligands by NEURL proteins

After establishing the differential activity of Neurl proteins, we wanted to test whether the NxxN motif was required for NEURL-mediated Notch activity in *Drosophila*. To do so, we exchanged one of the N’s in the NxxN consensus sequences of the ICDs of DLL1 and JAG1 to A or D (Fig. 4A-A’’’, Fig. 4S1). For Dl-DLL1, we chose to exchange the second N in the NBM motif to D to match the cryptic sequence in the DLL4-ICD (as discussed below). For Dl-JAG1, we chose to exchange the first N to A, to avoid disrupting an NN motif on the C-terminal of the NxxN motif, which has been shown to be weakly involved in deltaD signalling in zebrafish^29^. Both variants activate Notch in wing discs that express WT Mib1 (Fig. 4B-B’’). However, in contrast to Dl-DLL1 and Dl-JAG1, the activation of Dl-DLL1-N2D and Dl-JAG1-N2A was diminished upon co-expression with Neurl-1 in *mib1* mutant discs (Fig. 4C-C’). Moreover, Dl-rDll1 variants with the complete NxxN motif mutated to A or only the C-terminal N to A or D, could not be activated by Neurl-1 (D-rDll1-NBM2A, Dl-rDll1N2A and Dl-rDll1-N2D, Fig. 4S1 A-A’’). Similarly, Neurl-1 failed to activate Dl-Jag1 variants where either the NxxN motif is completely mutated, or only the N-terminal N was mutated (Dl-Jag1-NBM2A and Dl-Jag1-N2A, Fig. 4S1B, B’). These results indicate that the NxxN motif is required for the activation of DLL1 and JAG1 orthologs by NEURL-family members and that it is a functional NBM.

**Fig. 4.**
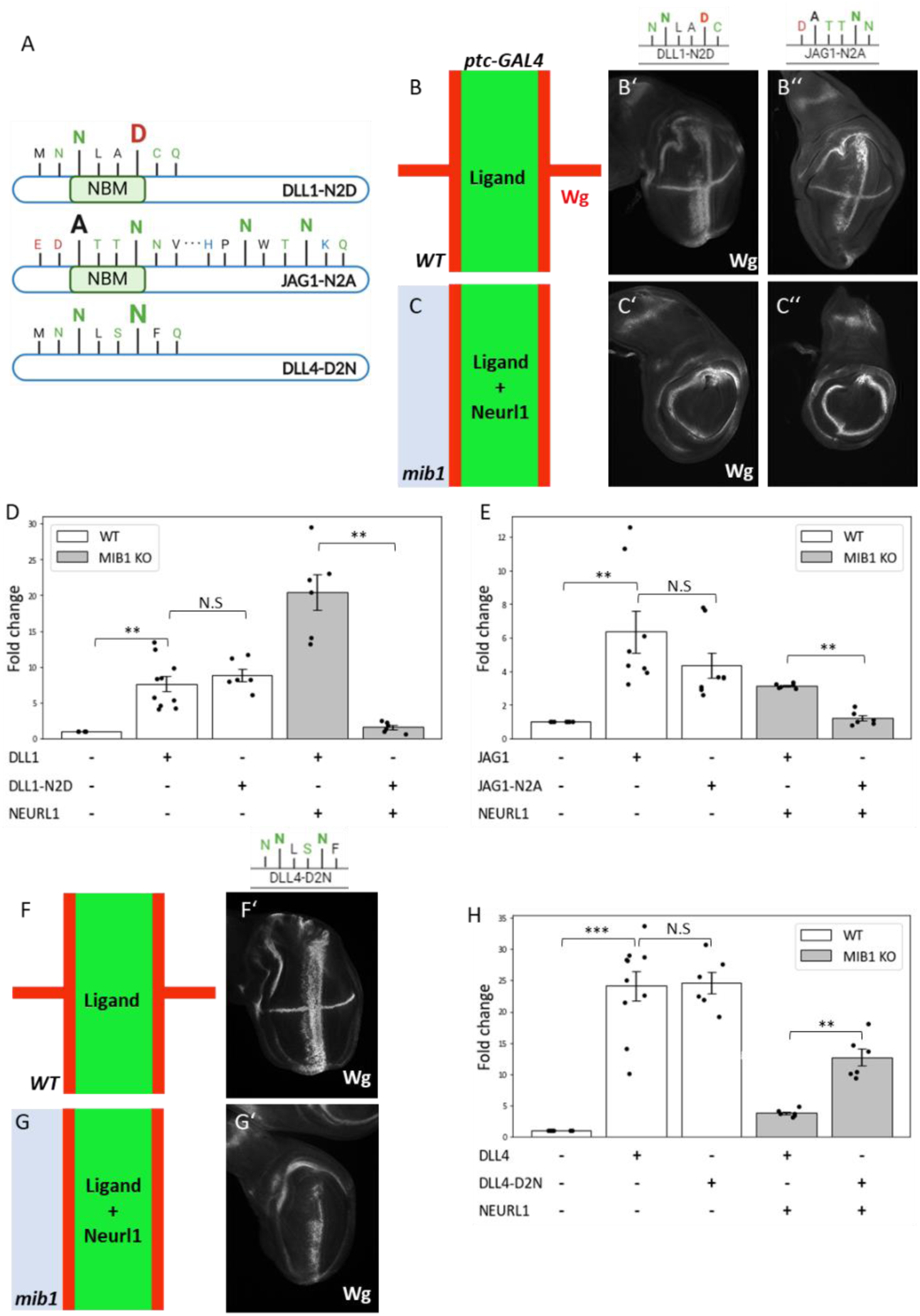
The NxxN motif is required and sufficient for activation of Notch ligands by NEURLs. (A) Schematic of the variant ICDs used in the experiment, with specific substitutions abolishing the NxxN motif in DLL1 and JAG1 ICDs, while reconstituting it in DLL4. (B-B’’) Expression of the mutated Dl-DLL1-N2D and Dl-JAG1-N2A in WT wing imaginal discs induces ectopic Wg expression, similar to unmutated Dl-DLL1 and Dl-JAG1. (C-C’’) Co-expression of these mutated hybrid ligands with Neurl-1 in mib1 mutant discs failed to activate Wg. (D-E) Luciferase activity assay comparing WT cells containing MIB1 and MIB1-KO cells, both co-expressing DLL1-N2D (D) or JAG1-N2A (E), co-cultured with Notch receiver cells. (F-F’, G-G’) Expression of the reconstituted Dl-DLL4-D2N in both WT wing (F-F’) and in Mib1 mutant wing when co-expressed with Neurl-1 (G-G’) induces ectopic Wg expression. (H) Luciferase activity assay comparing WT cells containing Mib1 expressing either non-mutated DLL4 or DLL4-D2N with Mib1-KO cells co-expressing NEURL-1 with either non-mutated Dll4 or DLL4-D2N, all co-cultured with Notch receiver cells. Results represent fold change compared to the negative control, with p-values measured by Mann-Whitney test (* - p<0.05, ** - p<0.01, ** - p<0.001, n>=3).

To further test the requirement for the NxxN sequence in DLL1 and JAG1 as MBM, we analysed the activity of NxxN variants in a cell culture assay. In this assay, we used sender cells expressing full-length ligands containing the same substitutions as above, namely, DLL1-N2D and JAG1-N2A and compared them to WT ligands. Luciferase activity assay showed that these variants can activate Notch in WT cells (containing MIB1) but cannot activate Notch in MIB1-KO cells expressing NEURL-1 (Fig. 4D, E). These results show that full-length DLL1 and JAG1 require the NxxN motif to be activated by NEURL-1. Thus, it is likely that these motifs are functional NBMs, as found for the binding partners of Neur in *Drosophila*.

The sequence comparison revealed that a cryptic NBM similar to that of DLL1 can be recognized in DLL4 with the C-terminal N exchanged by aspartic acid (D) (NLS<u>D</u> instead of NLA<u>N</u>, Fig. 4A). This exchange to D is conserved in all mammals and in chicken (Fig. 1S1A). In zebrafish, an exchange to the similar glutamic acid (E) can be observed. To test whether the exchange of N to D is the basis for the difference in the ability to be activated by NEURL proteins, we mutated the C-terminal D in DLL4 to N (NLS<u>N)</u>. Expression of Dl-DLL4-D2N in wt discs resulted in the induction of ectopic expression of Wg comparable to Dl-DLL4, indicating that Mib1-mediated signalling was not affected by the D2N mutation (Fig 4F, F’). However, in contrast to Dl-DLL4, Dl-DLL4-D2N was also able to activate Notch in *mib1* mutant discs upon co-expression with Neurl-1, indicating that it is the lack of the second N in this cryptic NBM that prevents the activation of DLL4 by Neurl-1 (Fig. 4G, G’).

To test whether the NxxN motif is sufficient for activation also in mammalian cells, we performed a cell culture assay with full length DLL4-D2N and compared its activity to full length DLL4 (Fig 4H). We found that both DLL4 variants similarly activated receiver cells when expressed in WT U2OS (containing MIB1). However, when co-expressed with NEURL-1 in MIB1-KO cells, DLL4-D2N showed a significantly higher activation compared to DLL4. This suggests that the presence of a NxxN motif is sufficient for activation in *Drosophila* and mammalian cells.

### Reduced co-localization between NEURL-1 and Notch ligands is observed when the NxxN motif is mutated

To verify that the discovered NBM is crucial for the interaction of NEURL-1 with Notch ligands, we performed a co-localization assay using fluorescent markers fused to the C-terminal of the ligands (mTurquoise2, mTQ2) and NEURL-1 (mCherry) to track their co-localization (Fig. 5A). As a measure of co-localization, we defined a co-localization ratio as the fraction of mCherry (fused to NEURL-1) pixels that co-localized with mTQ2 (fused to ligands) pixels. Analysis of the images (Figure 5B-D) showed that NEURL-1, when co-expressed with DLL1 or JAG1, was mainly distributed in vesicles, highly co-localized with the two ligands. In contrast, when NEURL-1 was expressed with Dll4, it was more evenly distributed in the cytoplasm with significantly less co-localization with the ligands. Moreover, we observed a significant decrease in co-localization between NEURL-1 and the two ligands, DLL1 and JAG1, when the NBM was mutated, implying that the NBM is required for the higher co-localization (Fig. 5B-C’). Finally, looking at the DLL4 variant in which the NBM has been re-introduced (DLL4-D2N) we found a significant increase in its co-localization with NEURL-1 (Fig. 5D, D’). These results confirm that the NBM is required and sufficient for the interaction of NEURL-1 with the Notch ligands.

**FIG. 5.**
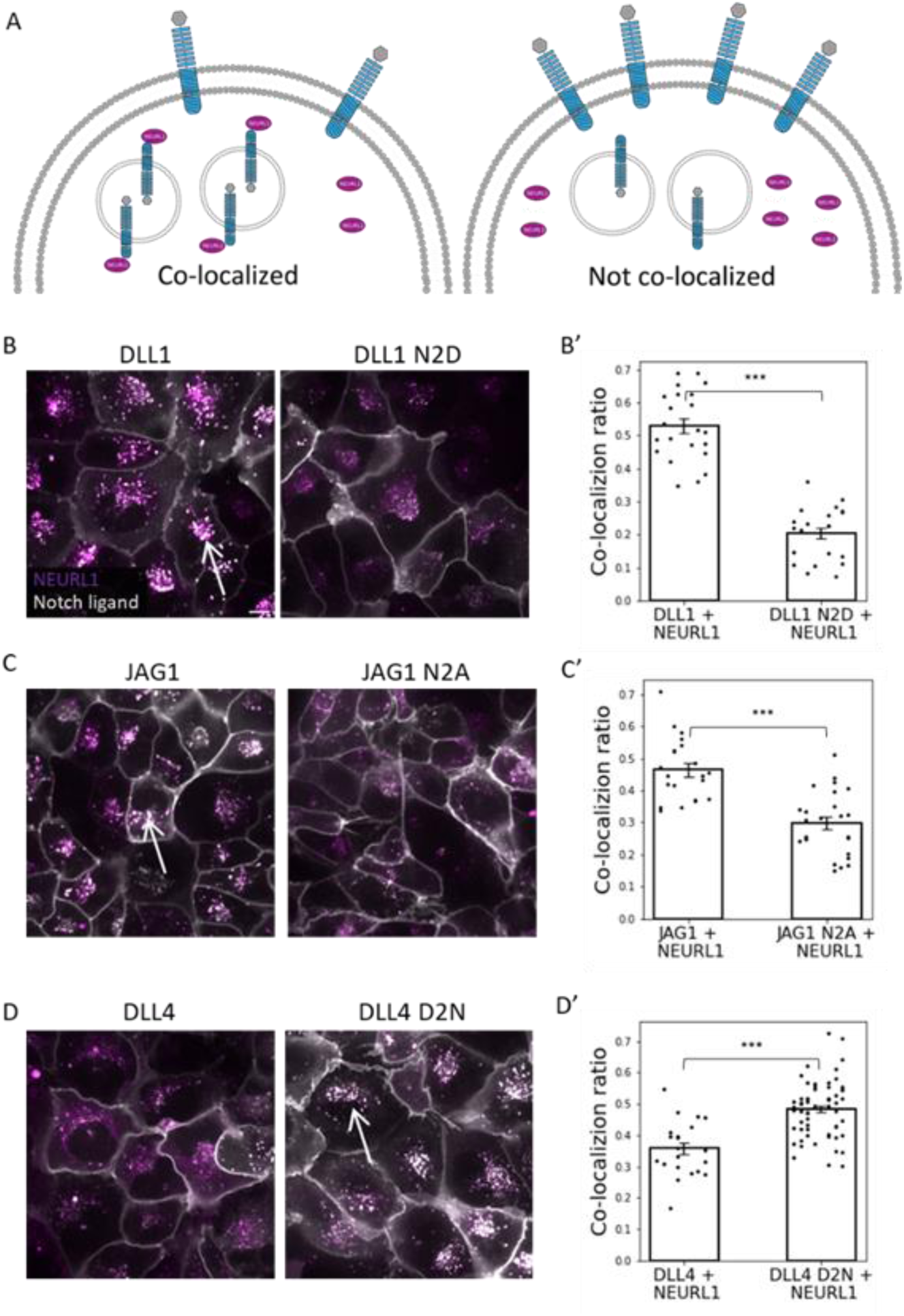
Enhanced co-localisation of NEURL-1 with Ligands depends on the presence of the NBM. (A) Co-localization scheme. Both NEURL-1 and the Notch ligand are marked with fluorescent proteins. This assay measures the co-localization ratio which is defined by the fraction of mCherry (fused to NEURL-1, shown as magenta) pixels that are co-localized wit mTQ2 (fused to ligands, shown as gray) pixels. (B-D) Representative images of cells co-expressing NEURL-1 fused to mCherry and one of the WT (left) or mutated (right) Notch ligands (as indicated) fused to mTQ2. (B’-D’) Quantification of co-localization of different Notch ligands (as indicated) with NEURL-1. P-values were measured by Mann-Whitney test (*** - p<0.005, n>=19 images per ligand / variant). Arrows mark examples of a co-localization spots.

### Analysis of the functionality of the mammalian ICDs in Neur-dependent signalling processes *in vivo*

We next asked whether the ICDs of DLL1, DLL4, JAG1, and JAG2 can complement the activity of Dl at the organismal level, particularly in well-characterized Neur-dependent developmental processes in *Drosophila*. For this purpose, we generated knock-in alleles encoding hybrid ligands in which the ICD of Dl is replaced by that of the mammalian ligands (Fig. 6). We utilized the Dl^attP^-landing site, as previously described in^30^. *Dl^attP^* is a null allele of Dl, causing embryonic lethality in homozygosity, because exon 6, which encodes most of Dl, is replaced by an attP landing site^31^. The mutant animals die as embryos due to hyperplasia of the nervous system, a neurogenic phenotype characteristic of Notch pathway mutants, also *neur* mutants^32^. This neurogenic phenotype is revealed by the reduction of the cuticle to a small dorsal patch in cuticle preparations (Fig. 6F, G).

**Fig. 6.**
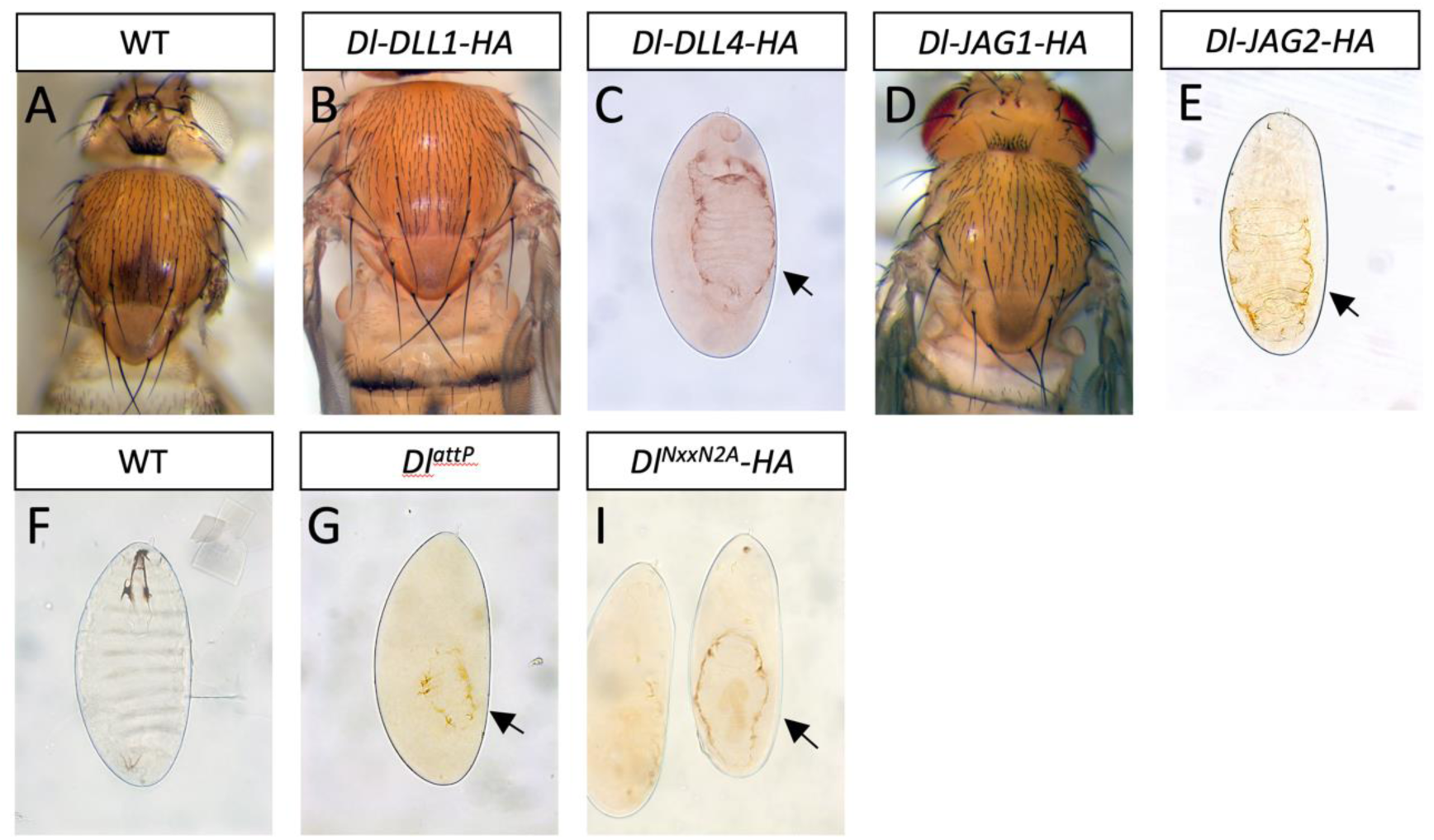
*In vivo* analysis of the ICDs of the mammalian ligands in *Drosophila*. For this purpose, knock-in alleles were generated that expressed the hybrid ligands instead of Dl. The phenotypes of flies homozygous for the alleles are shown. (A, B, D) Only the Dl-Dll1 and Dl-JAG1 flies develop to adulthood and therefore rescue the embryonic lethality. (C, E) In contrast, Dl-DLL4 and Dl-Dl-JAG2, which do not possess a NBM, die as embryos and display the typical neurogenic cuticle phenotype, also observed for homozygous *Dl^attP^* flies (F, G) and *Dl^NxxN2A^*, a Dl allele where the NBM is mutated to alanine (I, compare with F, G). The arrows in (C, E, G, I) point to the residual cuticle fragment, a phenotype that is typical for mutant alleles of genes that encode components of the Notch pathway (neurogenic phenotype). Compare with the cuticle preparation of a wt embryo, shown in (F).

We previously showed that the knock-in of the ICD of Dl into *Dl^attP^* resulted in a wildtype allele (*Dl^attP^-Dl-HA*) that provided full activity and allowed correct development of the flies. In contrast, the knock-in of the ICD of a Dl variant with a mutated NBM failed to rescue the neurogenic phenotype in homozygosity (*Dl^NxxN2A^*, Fig. 6F, I, arrow)^30^.

We found that, similar to *Dl^attP^-Dl-HA,* homozygous for *Dl^attP^-DLL1-HA* or *Dl^attP^-JAG1-HA* developed to the adult stage, indicating that the ICDs of DLL1 and JAG1 provide sufficient activity for the completion of development (Fig. 6A, B, D). In contrast, flies homozygous for *Dl^attP^-DLL4-HA* or *Dl^attP^-JAG2-HA* failed to rescue the *Dl*-mutant phenotype and died as embryos. The homozygous embryos displayed the typical neurogenic phenotype, indicating a failure of Neur-mediated activation (Fig. 6C, E, compare with G). These findings confirm that only ICDs containing an NBM can mediate DSL-signalling in Neur-dependent processes and demonstrate that the NBM of DLL1 and JAG1 is fully functional *in vivo*.

## Discussion

During *Drosophila* development, Neur-induced Notch-signalling plays a crucial role in establishing the nervous system by selecting neural precursor cells of the peripheral and the neuroblast of the central nervous system from an equivalence group^33–37^. In contrast, Neurl-1 and Neurl-2 double mutant mice develop normally without detectable developmental defects, neuronal or otherwise, indicating that the developmental roles of Neur have been lost in the transition between flies and mammals^38^. It appears that, in mammals, the developmental role of Neur has been taken over by MIB1 and it is unclear whether Neurl proteins are involved in Notch signalling in mammals at all. However, Neurl proteins do function in the adult mouse brain, where they are involved in neuronal plasticity and spatial memory, processes also dependent on Notch signalling^13,39^. Moreover, cell culture experiments show that Neurl-1 can ubiquitylate Jag1 and Neurl-2 can ubiquitylate Dll1, suggesting that Neurl proteins can bind to and ubiquitylate these ligands^10,40^. Interestingly, these *in vitro* experiments also suggest that the role of Neurl-mediated ubiquitylation in vertebrates is opposite to that in *Drosophila*, as it appears to down-regulate signalling by Jag1 in mammals and XDelta1 in Xenopus^10,40,41^.

Here we show, both *in vivo* (humanized *Drosophila*) and in mammalian cell culture, that NEURL-1 and NEURL-2, like Neur, can differentially activate DLL1 and JAG1 but not DLL4 and JAG2. This strongly suggests that both NEURL proteins are activators of DSL-signalling, acting similarly to Neur in *Drosophila*. The activation of the ligands by the NEURL proteins is dependent on the presence of a NxxN sequence in the N-terminal region of their ICDs. This consensus sequence, previously identified as NBM in *Drosophila*, is present in all investigated mammalian DLL1 and JAG1 homologs, which can be activated by the NEURL proteins, but not in DLL4 and JAG2 homologs^7^. The same holds true for the orthologs of *Xenopus laevis* (Fig. 1S1). Thus, it correlates with the ability of a ligand to be activated by the NEURL proteins, suggesting its importance for their function. The presented data confirm this conclusion. Mutating one N of the NxxN consensus sequence of DLL1 and JAG1 suppressed the ability of Neur and NEURL-1 and −2 to activate these ligands in *Drosophila* and cell culture experiments. Moreover, the knock-in of the ICDs of the mammalian ligands in the *Dl* locus showed that only the ligands with the NxxN sequence, DLL1 and JAG1, can prevent the development of a neurogenic phenotype in the embryo, also typical for loss of function of *neur*. Thus, the NxxN sequences in DLL1 and JAG1 function as NBMs also *in vivo*.

An interesting evolutionary aspect is the presence of a cryptic NBM in mammalian Dll4, where the C-terminal N is replaced by a D. This replacement of N to the acidic D in DLL1 abolished the activation by the NEURLs. Our experiments indicate that DLL4 can be activated in Mib1-dependent processes as efficiently as DLL1, suggesting that, unlike DLL1 and JAG1, it is devoted to Mib1-dependent signalling. However, replacing D by N in the cryptic NBM of DLL4 enabled its activation by NEURLs, indicating that the presence of the NxxN motif is sufficient for NEURL-dependent activation. Our results also suggest that JAG2 has a cryptic NBM, as it contains a similar cryptic sequence, with one N replaced by a similar acidic amino acid, E. Also in this case, this exchange is absolutely conserved among the mammalian JAG2 orthologs investigated (Fig. 1S1).

Previous work tested the effect of Neurl proteins in cells and in Xenopus and obtained results that contrast with previous reports that conclude that Neurls are suppressors of Notch signalling in vertebrates^10,40,42^. We explain this discrepancy by the concentration dependent manner by which Neur acts as an activator. Since Neur family members are potent inducers of endocytosis of DSL ligands (Fig 5), balancing Dl levels at the cell surface and the endocytosis efficiency required for ligand activation is crucial. This balance likely depends on Neur expression levels, as over-expression experiments in *Drosophila* suggest^35^. Thus, strong over-expression of NEURL-1 or −2 might accelerate ligand endocytosis, reducing surface ligand levels available for signalling, thereby reducing signalling, especially as MIB1 is likely also present in the cells and tissue used for the experiments. Our experimental set up was different since we used mammalian cells lacking the function of MIB1. This enabled us to monitor NEURL protein’s activation ability in the absence of possible competition with MIB1, which also induces endocytosis. It is possible that the additional presence of Mib1 with NEURL in a cell results in a reduction of the Notch pathway’s net activity in comparison with the sole presence of Mib1 or Neurl.

It is interesting to note that the heterozygous mutants of Jag1 and Notch1, as well as concomitant loss of Neurl-1 and −2, all cause spatial memory deficits in adult mice^13,15^. Our results linking Neurl activity to specific ligands could suggest a Neurl-Jag1 axis that operates during spatial memory and cognitive functions. Additional experiments may be required to specifically associate Neurl activity with cognitive functions in mammals. Interestingly, Neur appears to be involved in long-term memory formation in adult flies, suggesting the post-embryonic function of Neurl family members in the brain is evolutionarily conserved and might be their initial function^43^.

Our study shows that all Dl-hybrids can activate the Notch pathway in *Drosophila* imaginal disc cells, where only Mib1 is present, except for late occurring single sensory organ precursor cells. This indicates that, similar to mammalian MIB1, *Drosophila* Mib1 can productively interact with the ICDs of all functional mammalian DSL ligands. Thus, the mechanism of Mib1-mediated activation of the Notch pathway is conserved, and its mammalian version can be studied in these ‘humanized *Drosophila’*, where countless techniques allow detailed analysis.

Overall, our results support a previously unidentified regulatory mechanism for differential activation of Notch signalling in mammals, based on the ubiquitylation of the ICDs of Notch ligands. Hence, despite the fact that all Notch ligands can interact with all Notch receptors, the activity is restricted to specific ligands based on post-translational modifications in the ligands’ ICD. It will be interesting to check if this mechanism is relevant for other signalling pathways that involve combinatorial interactions between receptors and ligands.

## Acknowledgements

We thank Stefan Kölzer for excellent support in the generation of the described transgenic flies. We thank F. Schweisguth and G. Boullianne for providing for providing fly stocks. We thank Steve Blacklow for providing the Notch-Gal4 and MIB1-KO cell lines.

## Methods

### Fly stocks

*mib1^EY09870^* (Lai EC, 2005), *ptc*GAL4 (Speicher SA, 1994), *Dl^attP^* (Viswanathan R, 2019), *Dl^attP^-Dl-NEQN2A-HA* (Troost, 2023), *Dl^attP^-hDLL1-HA, Dl^attP^-hDll4-HA, Dl^attP^-hJAG1-HA, Dl^attP^-hJAG2-HA*, UAS-Dl-rDll1-HA, UAS-Dl-rDll1-N587A-HA, UAS-Dl-rDll1-N587D-HA, UAS-Dl-rDll1-NBM2A-HA, UAS-Dl-hDLL1-HA, UAS-Dl-hDll1-N595D-HA, UAS-Dl-hDLL4-HA, UAS-Dl-hDLL4-D573D-HA, UAS-Dl-hJAG1-HA, UAS-Dl-hJAG1-N1110A-HA, UAS-Dl-hJAG2-HA, UAS-mNeurl1-myc, UAS-hNEURL2-myc, UAS-Dneur-V5 (this study).

### Antibody staining of wing imaginal discs

Wing imaginal discs of wandering L3 larvae were dissected in PBS, fixed with 4% paraformaldehyde in PBS (4%PFA) for 30 minutes and washed with 0.3% Triton X-100 in PBS (PBT). Permeabilization and blocking was done using and 5% normal goat serum (NGS) in PBT for 30 minutes on room temperature. Primary antibody incubation was performed in 5% NGS in 0.3% PBT for 2 hours on room temperature followed by three washing steps with PBT. The corresponding secondary antibody was applied in 5% NGS in 0.3% PBT for 2 hours on room temperature. The following antibodies were used: mouse anti-Wg 4D4 (1:10, Developmental Studies Hybridoma Bank (DSHB), Iowa City, IA, USA), rabbit anti-β-Gal (polyclonal) (1:5000, MP Biomedicals, LLC., Solon, Ohio). Fluorophore-conjugated secondary antibodies were purchased from Invitrogen. Nuclei staining was done using the Hoechst 33258 dye. For further information see Klein^44^.

### Generation of DNA constructs

The constructs mNeurl1 and hNEURL2 were cloned from pEGFP-Vector into pUAST-attB-Vector for GAL4-driven expression in *Drosophila* via restriction cloning with the enzymes EcoRI and XhoI for mNeurl1 and EcoRI and XbaI for hNEURL2 and tagged with a C-terminal myc epitope. The constructs were inserted into the landing site 86Fb on the third Chromosome and recombined with the allele mib1^EY09780^. The Hybrid Ligands Dl-rDll1-HA, Dl-hDLL4-HA, Dl-hJAG1-HA and Dl-hJAG2-HA were generated via Overhang extension PCR and cloned into the pUAST-attB-Vector via the restriction sites NotI and XbaI and inserted into the landing site 51C on the second Chromosome. The ICD of hDLL1 was ordered from IDT technologies and cloned into the pUAST-attB-Dl-HA vector via the restriction sites BstEII and XbaI. Single point mutations were introduced via SDM. For endogenous expression under the Delta Promotor the Hybrid-ligands were cloned into the pGE-attB-Vector with the restriction enzymes BstEII and XbaI and inserted into the *DlattP* landing site^31^.

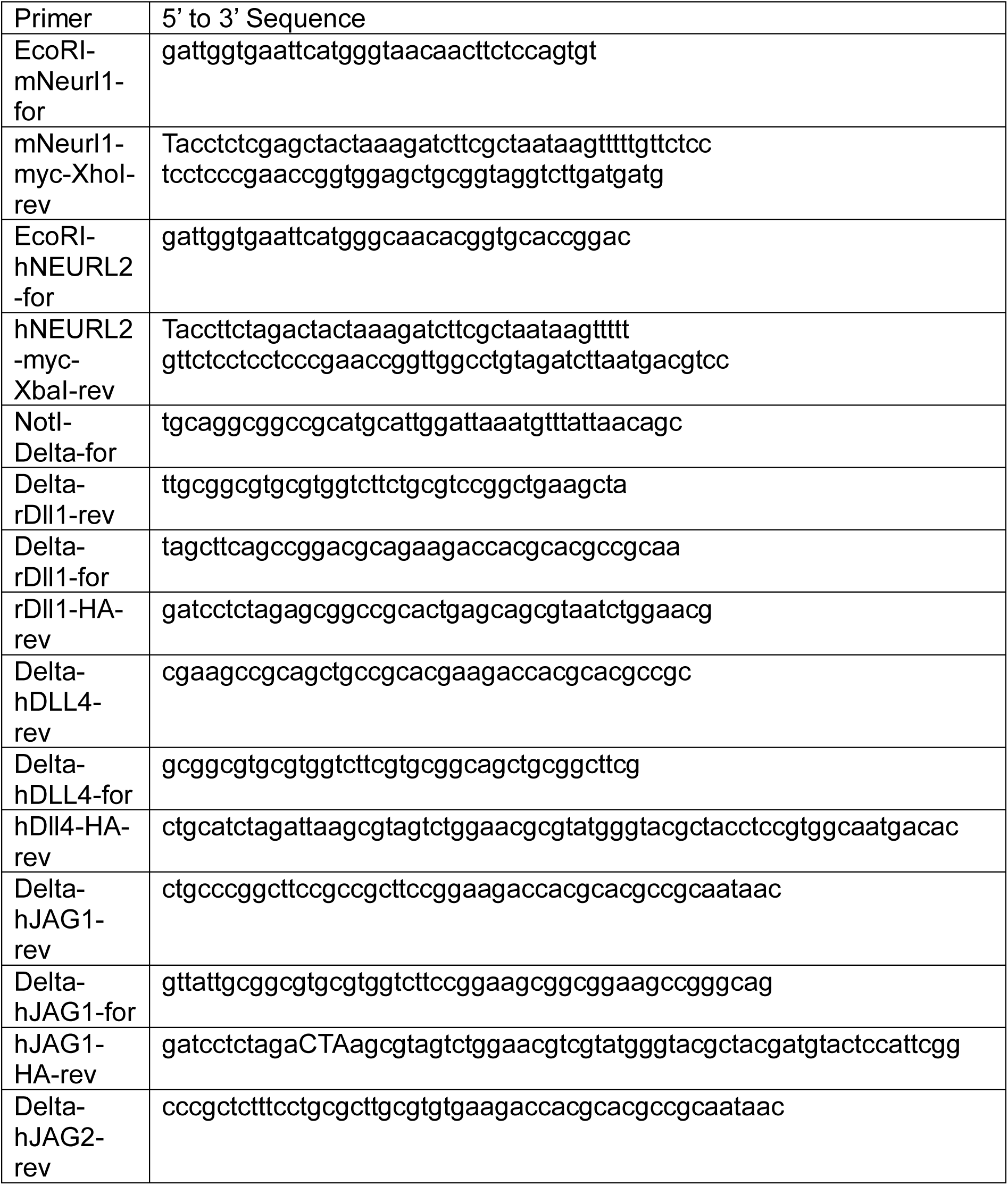
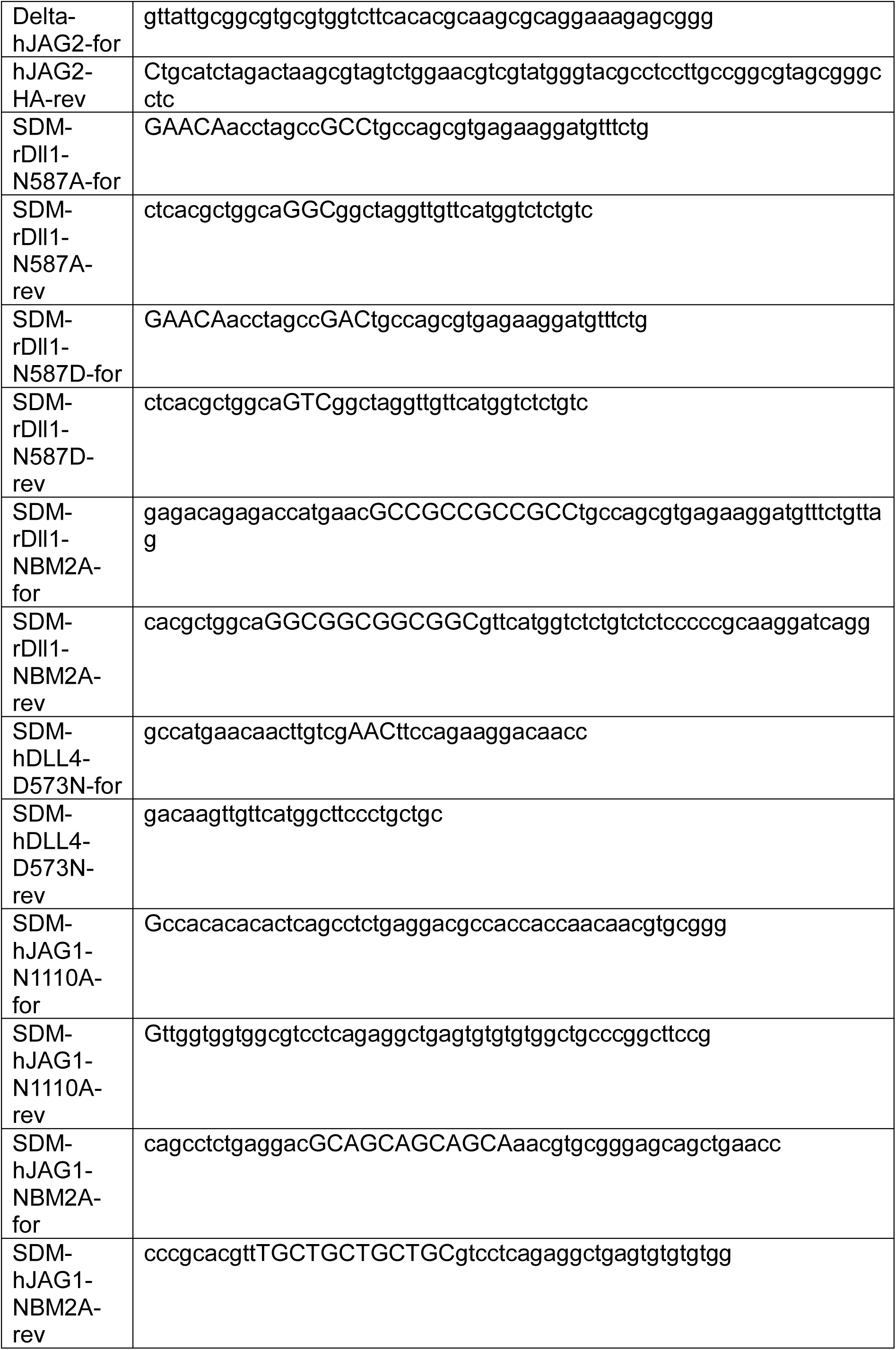
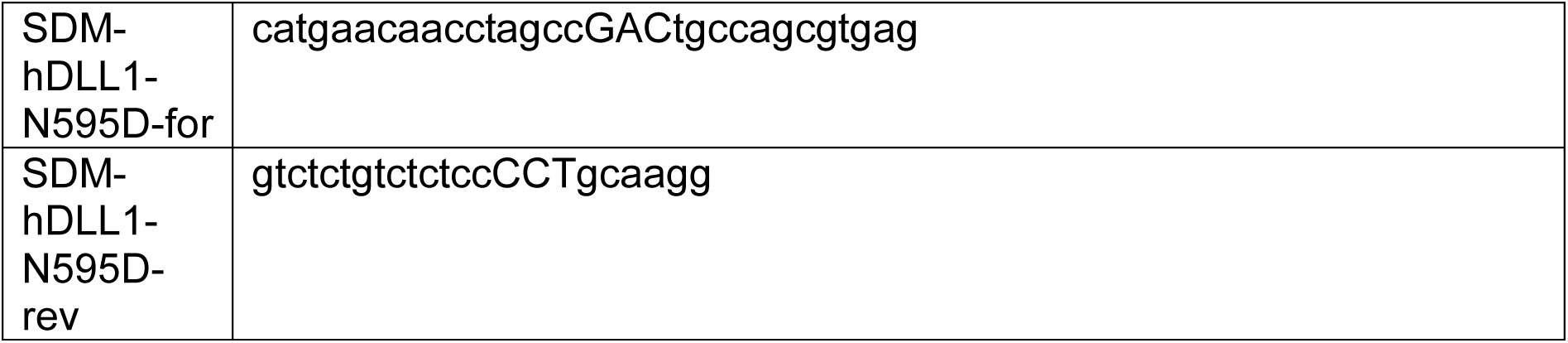

### Cells, materials and constructs

In this study, we used HEK293T (for lentivirus production) and U2OS cell lines, which were cultured in Dulbecco’s Modified Eagle’s Medium (DMEM) supplemented with 10% Fetal Bovine Serum (FBS). The cells were maintained at a constant temperature of 37°C and an atmosphere of 5% CO_2_. Transfection of the cells was executed using Lipofectamine 3000 (Thermo Fisher Scientific) following the manufacturer’s instructions.

Sender cells were generated through lentiviral transduction. This involved employing a pLVX vector encoding one of the ligands or NEURL-1, co-transfected with helper plasmids (pMD2.G and PsPax2), into HEK293T cells using Lipofectamine 3000. Following a 48-hour incubation period, the supernatant was carefully collected, filtered using a 0.45µm filter, and subsequently utilized for transduction on either U2OS WT or U2OS Mib1 KO cells.

Our receiver cells, U2OS-Notch1-Gal4 and U2OS-Notch2-Gal4 were kindly provided by Stephen C. Blacklow (Harvard Medical School)^27^.

### Luciferase Activity Test

The activity of NEURL-1 with the different ligands was measured with a luciferase activity assay. The detailed protocol for the luciferase assay is described in ^25^. Briefly, U2OS-NOTCH1-Gal4 or U2OS-NOTCH2-Gal4 (receiver cells) were cultured in a 24-well plate and transfected with 350 ng of UAS-firefly luciferase reporter^45^ and 10 ng of pRL-SV40 Renilla luciferase. Transfection was done using Lipofectamine 3000 (Thermo Fisher Scientific). 24h after transfection the receiver cells were transferred to a new 24-well plate with doxycycline (200ng/ml), co-cultured with sender cells expressing NEURL-1 and one of the Notch ligands. After additional 24h the cells were lysed with passive lysis buffer (Promega, 100µl/well) and taken to be measured with luminometer software (GloMax(R) Navigator Microplate Luminometer, Navigator 2010, Promega). Notch activity was defined as the ratio of luciferase to renilla, which was then normalized by the negative control to represent fold change in activity.

### Microscopy

Images of imaginal discs were acquired with the Zeiss Axio Imager Z1 Microscope equipped with a Zeiss Apotome or Apotome2.

Imaging of cells utilized an Andor revolution spinning disk confocal microscope, powered by 50 mW lasers from Andor in Belfast, Northern Ireland. The microscope featured a 37 °C temperature-controlled chamber and a CO_2_ regulator providing 5% CO_2_, both supplied by Okolab in Italy. The setup comprised an Olympus inverted microscope with an oil-immersion Plan-Apochromatic 60× objective NA = 1.42 from Olympus in Tokyo, Japan, along with an ANDOR iXon Ultra EMCCD camera also from Andor in Belfast, Northern Ireland. Control of the equipment was facilitated by Andor iQ software from Andor in Belfast, Northern Ireland.

### Co-localization assay

The co-localization assay was performed using live imaging microscopy with cells co-expressing NEURL-1 (fused to mCherry) and a DSL-ligand (fused to mTQ2). Snaps were taken in two channels, red and blue (561nm and 445nm respectively). All images were analyzed using custom code in python (https://github.com/Oren-Gozlan/Co-localization-code) to calculate the co-localization ratio. This was performed by taking all the pixels that have both red and blue values (above a threshold) and dividing them by all the red pixels (above the same red threshold). Each DSL-ligand was compared to its NBM related variant (e.g DLL1 vs DLL1-N2D) and significance between samples was calculated with a Mann Whitney U test.

## Supplementary Figure captions

***FIG. 1S1.** (A) Sequence comparison of the ICDs of vertebrate orthologs of Dll1, Dll4, Jag1 and Jag2. In Dll1 and Jag1, the NxxN motif is conserved (highlighted area), as well as neighbouring amino acids. In Dll4 and Jag2 one of the essential Asparagines is replaced with D or E (highlighted area). (B-B’’’) Expression of the Dl-hybrids in mib1 mutants fails to induce ectopic expression of Wg, indicating that their activity in the wing disc depends on Mib1*.

**Fig. 3S1.** Dl-rDLL1 hybrid recapitulates the results found with Dl-hDLL1. Wing discs expressing Dl-rDLL1 with ptc-Gal4 in WT (A), mib1 mutant(A’), and mib1 mutant with over-expression of Dneur (A’’)/ Neurl-1 (A’’’)/ NEURL-2 (A’’’’).

**Fig. 4S1.** The NxxN motif in Dl-rDll1 is a NBM that is required for activation of the ICD by Neurl-1. (A-B’) Wing discs from mib1 mutant flies expressing Neurl-1 and variants of Dl-rDLL1 or Dl-JAG1 (top labels) in which the entire NBM is replaced by Alanines (NBM2A) (A and B) or only one Asparagine is replaced by Alanine or Aspartic Acid (N2A, N2D) (A’, A’’ and B’). All variants without a complete NBM failed to activate Wg.

