## Supplemental figures for "Neuralized-like proteins differentially activate Notch ligands"

Supplementary Figure 1S1

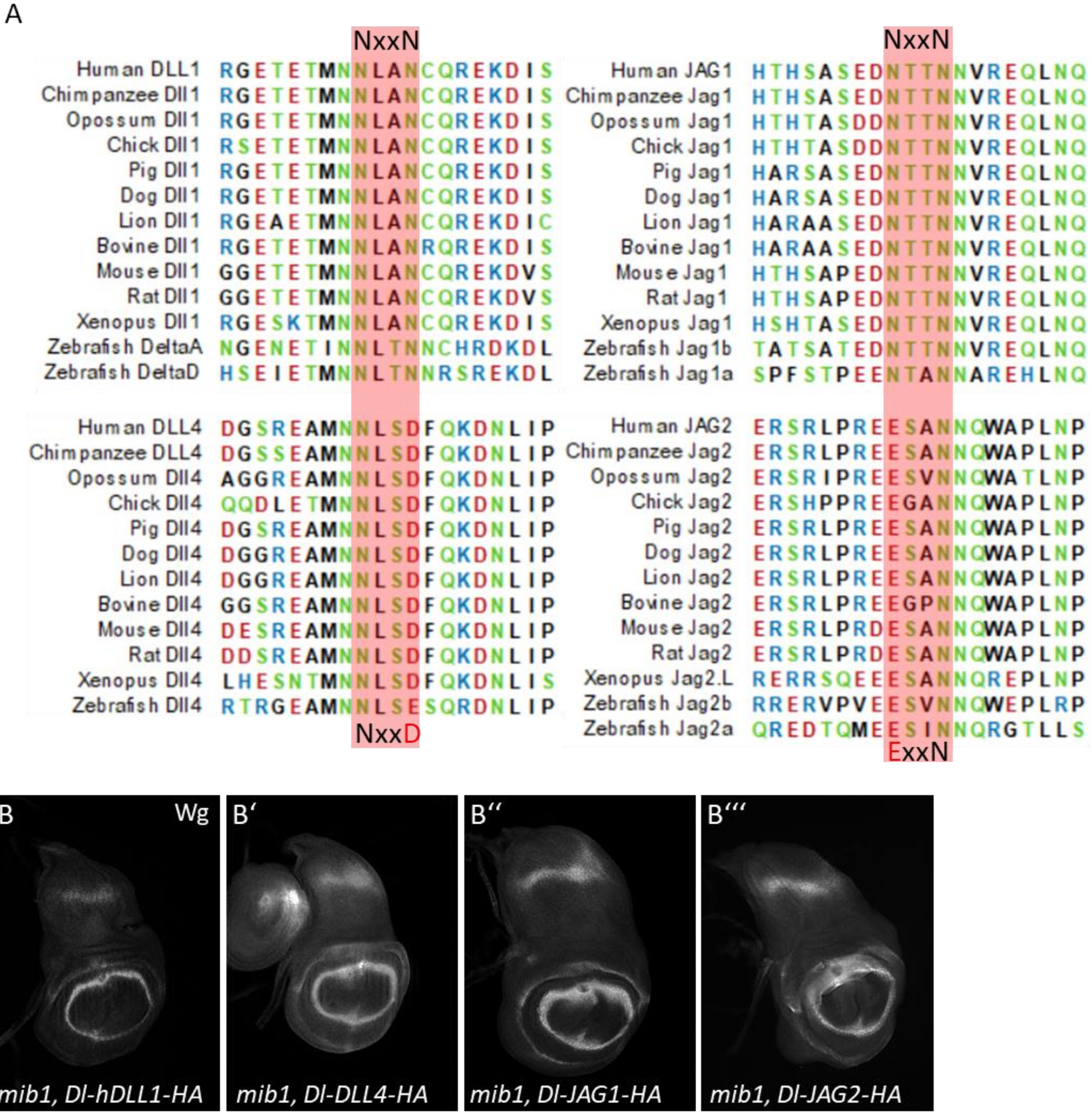


***FIG. 1S1***. *(A) Sequence comparison of the ICDs of vertebrate orthologs of Dll1, Dll4, Jag1 and Jag2. In Dll1 and Jag1, the NxxN motif is conserved (highlighted area), as well as neighbouring amino acids. In Dll4 and Jag2 one of the essential Asparagines is replaced with D or E (highlighted area). (B-B’’’) Expression of the Dl-hybrids in mib1 mutants fails to induce ectopic expression of Wg, indicating that their activity in the wing disc depends on Mib1.*

Supplementary Figure 3S1

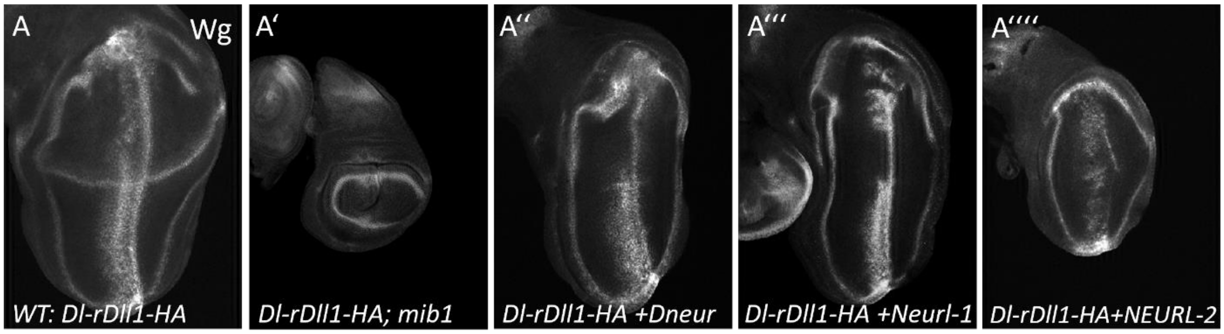


**Fig. 3S1**. *Dl-rDLL1 hybrid recapitulates the results found with Dl-hDLL1. Wing discs expressing Dl-rDLL1 with ptc-Gal4 in WT (A), mib1 mutant(A'), and mib1 mutant with over-expression of Dneur (A'')/ Neurl-1 (A''')/ NEURL-2 (A'''').*

Supplementary Figure 4S1

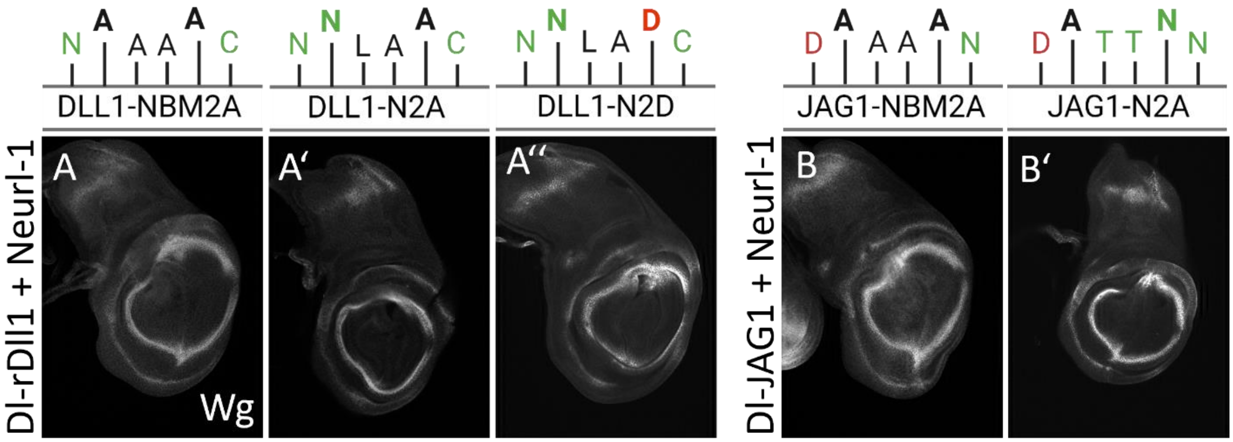


**Fig. 4S1**. *The NxxN motif in Dl-rDll1 is a NBM that is required for activation of the ICD by Neurl-1. (A-B') Wing discs from mib1 mutant flies expressing Neurl-1 and variants of Dl-rDLL1 or Dl-JAG1 (top labels) in which the entire NBM is replaced by Alanines (NBM2A) (A and B) or only one Asparagine is replaced by Alanine or Aspartic Acid (N2A, N2D) (A', A'' and B'). All variants without a complete NBM failed to activate Wg.*
